# Deriving functional network topology from *in vivo* two-photon calcium imaging: state-dependent graph features in mouse mesoscale motor cortical network

**DOI:** 10.64898/2026.03.27.714836

**Authors:** Genchang Peng, Nurullah Sati, Shahrzad Latifi

**Affiliations:** Department of Neuroscience, Rockefeller Neuroscience Institute, West Virginia University, Morgantown, WV, USA

**Keywords:** Mesoscale networks, graph-theoretical analysis, primary motor cortex, two-photon calcium imaging, functional connectivity, small-world topology, hub organization

## Abstract

Mesoscale neuronal networks represent an intermediate organizational level linking single-neuron activity to large-scale brain networks. Here, we used in vivo two-photon calcium imaging and graph-theoretical analysis to characterize functional network topology in the primary motor cortex across behavioral states. Motion networks exhibited the largest functional connectivity architectures, whereas anesthesia networks showed reduced network scales together with stronger modular segregation and more pronounced small-world topology. Network sign further shaped topology, with negative associations associated with reduced modularity and weakened small-world structure. Hub analyses revealed additional state-dependent differences: anesthesia networks exhibited stronger hub connectivity despite reduced neuronal activity, whereas motion networks showed higher hub activity with weaker connectivity structure. These findings demonstrate that mesoscale neuronal networks exhibit structured and state-dependent organization and provide a framework for studying cortical network dynamics in normal brain function and brain disorders.

## Introduction

The brain is a complex system of highly optimized interconnected networks, organized across multiple spatiotemporal scales^1^. Complex systems, despite differences in nature, scope, or governing mechanisms, exhibit convergent organizational principles, such as modular structure, hierarchical structuring, and small-world topology^2, 3^. Whitin this context, brain networks share emergent features characteristic of complex adaptive systems. These organizational attributes support the balance between functional segregation and integration in the brain, enabling efficient information transfer across distributed multiscale networks encompassing genetic, molecular, neuronal, circuits and regional levels^4^.

The emergent organizational characteristics of brain networks have been extensively investigated to elucidate mechanisms underlying normal brain function and to characterize network alterations associated with neurological disorders^5, 6^. To date, the majority of these studies have relied on data derived from large-scale (macroscale) brain networks, primarily using neuroimaging and electrophysiological modalities such as functional magnetic resonance imaging (fMRI) and electroencephalography (EEG) ^1, 7, 8^. While these approaches have provided foundational insights into systems-level connectivity, their spatial resolution limits interrogation of network dynamics at the cellular and microcircuit levels.

Advances in brain monitoring technologies over the past decade have enabled functional network interrogation at the mesoscale, bridging the gap between microscale neuronal activity and/with macroscale systems organization. Mesoscale brain networks are defined as the intermediate organizational level linking individual cells to local ensembles and functional circuits, capturing emergent connectivity patterns that underlie circuit-level information processing ^1, 9-11^. Among advanced monitoring approaches, in vivo two-photon (2P) calcium imaging offers longitudinal tracking of mesoscale brain network with cellular resolution during behavior in both physiological and pathological conditions^12^. When integrated with graph-theoretical frameworks, mesoscale imaging data provide a powerful platform for quantifying functional network topology. Network features, including functional connectivity, clustering architecture, path efficiency, and modular organization, can be systematically evaluated to characterize communication structure within neuronal populations^13-17^. However, despite increasing application of mesoscale imaging approaches, comprehensive characterization of higher order network properties remains incompletely understood at this scale.

In this work, we leveraged in vivo 2P calcium imaging to interrogate excitatory cortical neuronal networks at the mesoscale level. Pairwise Pearson correlations (PC) were computed to construct singed functional connectivity (FC) matrices, and graph theoretical metrics were applied to quantify global and nodal topology. By systematically comparing motion, no-motion, and anesthesia conditions, we characterized state-dependent variations in network scale, edge composition, small-world organization, modular structure, and hub characteristics (architecture). Our data demonstrates that higher-order network features can be robustly extracted from mesoscale calcium imaging data using a validated signed connectivity framework. Together, this approach provides a multiscale network perspective on how mesoscale cortical connectivity reorganizes across distinct brain states.

## Results

### 1. State-dependent connectivity analysis

To assess neuronal population activity in the primary motor cortex (M1), a CaMKII-tTA/tetO-GCaMP6s transgenic mouse line was used, enabling stable and uniform expression of the genetically encoded calcium indicator GCaMP6s in excitatory neurons^12^. A 5-mm cranial window was stereotaxically implanted over the unilateral forelimb region of M1 in adult male mice, allowing in vivo two-photon calcium imaging of Layer II/III neuronal ensembles during behavior. A motorized wheel was used to induce motion at constant velocity for the motion condition, while an anesthesia apparatus was used to establish the anesthesia state. 2P imaging datasets were subsequently processed using the Suite2p analytical pipeline to extract ΔF/F activity traces derived from fluorescence signals^18^. To mitigate slow fluorescence decay of GcaMP6s and sharpen the calcium transient, deconvolution was applied to Δ*F/F activity traces*, yielding deconvoluted signals used for subsequent PC calculations (Fig.1)^19-21^. Since PC preserves signs, this quantitative approach produced both positive and negative associations in the raw correlation matrices (Fig.1b). Positive correlations might reflect synchronous activity or coordinated ensemble dynamics among neuronal populations^22-24^, whereas negative correlations could represent anti-correlated or antagonistic interactions^25-27^. Given the large number of recorded neurons, raw PC matrices were inherently dense and contained spurious correlations arising from spatial proximity, data noise or global signal fluctuations. To address this limitation, correlation estimates were subjected to statistical validation before being interpreted as FC. Accordingly, surrogate testing and multiple comparison correction were adopted as validation strategy to identify significant correlations and construct statistically supported FC networks. Surrogate testing was performed by independently shuffling the deconvoluted Δ*F/F* activity traces for each neuron (*K* = 1000 iterations) to generate null distribution of correlation values^28^. Edges exceeding the 95% confidence interval (*p <* 0.05) were retained. Significant associations were subsequently corrected using Benjamini-Hochberg procedure to control the false discovery rate (FDR; *q* = 0.05)^29^. Following this two-step validation framework, the dense raw correlation matrices were transformed into sparse, statistically supported functional connectivity architectures (Fig.1d).

Application of the two-step validation framework substantially reduced network density relative to raw PC matrices. As shown in Figure 2a, these reductions were highly significant across all three states (no-motion: *p* = 6.39 × 10^−9^, *n* = 5; motion: *p* = 2.15 × 10^−9^, *n* = 5; anesthesia: *p* = 1.70 × 10^−5^, *n* = 3; paired *t*-test), confirming that only a subset of correlations represented statistically reliable functional interactions. The resulting validated FC networks exhibited clear state-dependent differences in organizational scales. Motion networks demonstrated the largest architecture, characterized by the highest number of nodes (307.21 ± 96.70) and significant edges (3504.22 ± 1736.15), whereas anesthesia networks showed the most reduced connectivity structure (node: 245.33 ± 64.82 and edge:1662.01 ± 290.15 counts). No-motion networks served as an intermediate baseline configuration. Statistical comparison confirmed significant differences between motion and anesthesia states in both nodal representation and edge density, while no-motion did not differ significantly from either condition (*p >* 0.05, unpaired *t*-test). Collectively, these findings position motion and anesthesia as the two extremes of mesoscale network organization, reflecting expanded (motion) versus constrained connectivity architectures (anesthesia), while no-motion representing a transitional baseline state.

**Figure 1.**
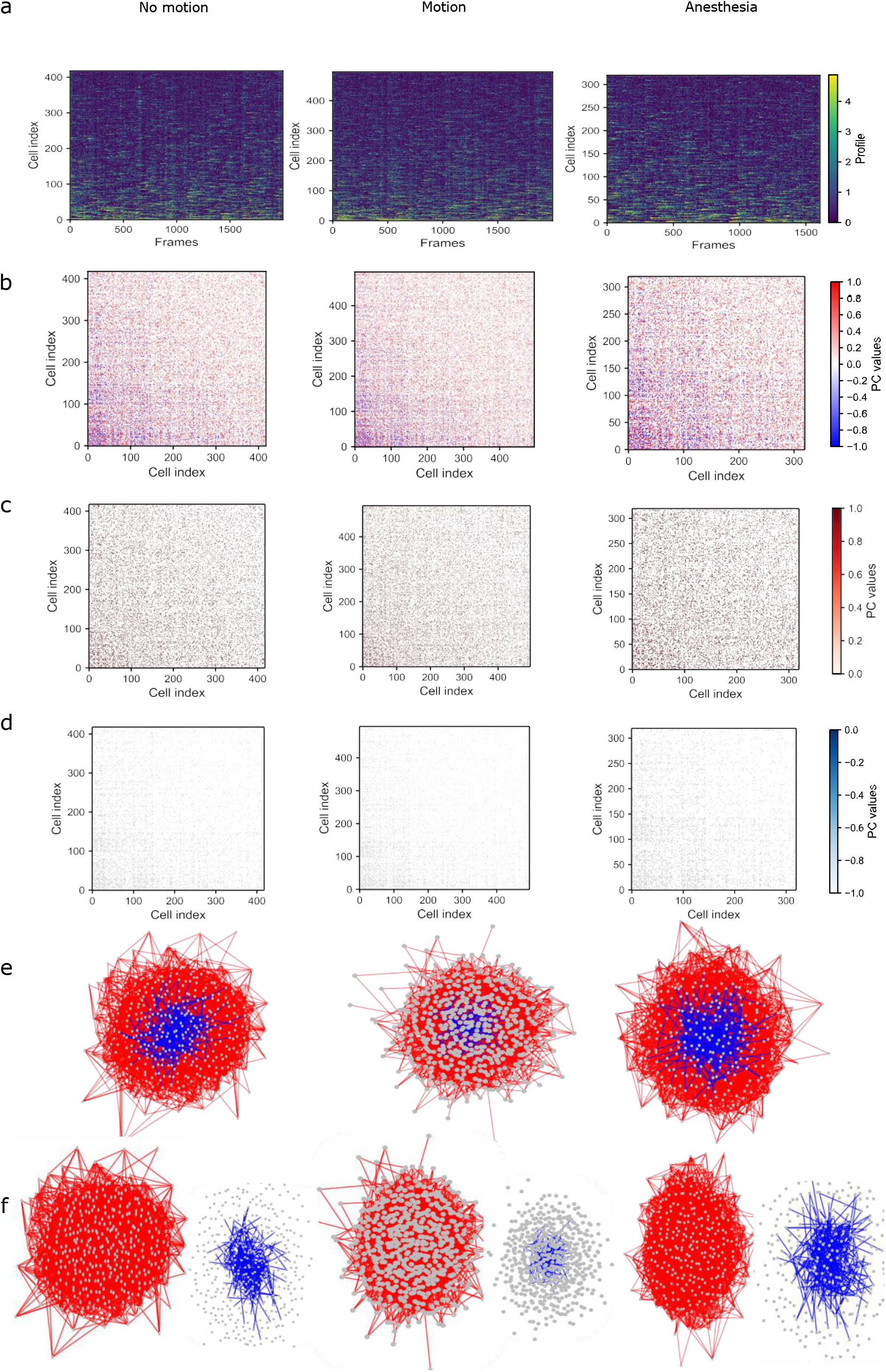
Functional correlation patterns across behavioral states. (a) Deconvolved Δ*F/F* neuronal traces of three states. (b) Corresponding Pearson correlation (PC) matrices. (c) positive PC matrices. (d) negative PC matrices. Left to right: no motion, motion and anesthesia. (e) Graph visualizations of functional connectivity (FC) networks. Positive and negative edges are separated in red and blue colors.

**Figure 2.**
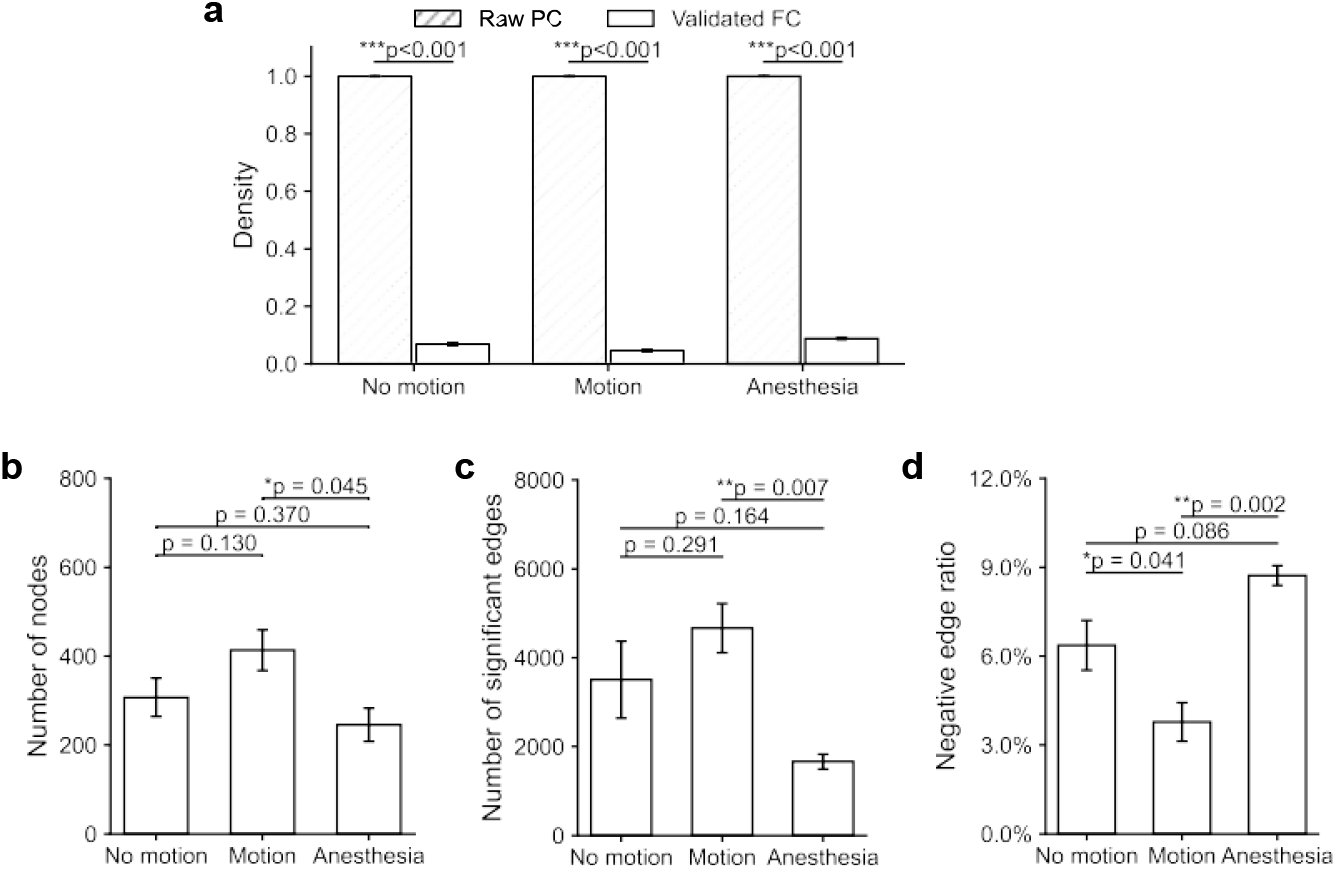
State-dependent reorganization of functional connectivity (FC) network architectures. (a) Network density is significantly reduced in validated FC networks compared to their respective raw Pearson correlation (PC) matrices across all three states. Paired t-test, *n* = 5 for motion, *n* = 5 for no motion, *n* = 3 for anesthesia. (b, c) Comparison of network nodes and number of edges in the validated FC networks among three states. Data points represent individual recordings; *p*-values denote unpaired *t*-tests. (d) Proportions of negative edges show significant elevations in anesthesia compared to no motion and motion. Error bars represent SEM. * *p <* 0.05, ^**^*p <* 0.01, ^***^*p <* 0.001.

Across all states, validated FC networks comprised both positive and negative correlations, reflecting the coexistence of both synchronized and antagonistic neuronal interactions^26, 30^. However, negative edges constituted only a minor fraction of overall connectivity with mean proportions remaining below 10% across conditions. Motion networks exhibited the lowest fractions of negative edges, whereas anesthesia showed the highest (no-motion: 6.42 ± 1.70%; motion: 3.84 ± 1.22%; anesthesia: 8.78 ± 0.56%; Fig. 2d). Statistical comparisons confirmed a significant elevation of negative edge proportions during anesthesia relative to both non-motion baseline and motion states (p <0.05). These findings indicate that negative functional associations are state-dependent and differentially expressed across brain states.

In summary, application of the two-step validation framework transformed dense raw correlation matrices into sparse, statistically supported FC networks derived from deconvoluted ΔF/F activity traces. Network scale exhibited clear state dependence, with motion and anesthesia representing the largest and smallest connectivity architectures, respectively, and no-motion occupying an intermediate baseline configuration. Negative edges comprised a minor but state-dependent component of network organization, remaining below 10% across conditions and highest under anesthesia.

### 1. Network topology metrics

To characterize global network organization, we quantified graph-theoretical properties of validated functional connectivity networks across states and network types. These metrics capture complementary aspects of network topology, including modular organization, small-world structure, and hub-centered architecture.

### 1.1 Modularity

Modularity quantifies the extent to which a network can be partitioned into distinct communities characterized by dense intra-community connectivity and comparatively sparse connections between communities^31, 32^. In brain networks, modular organization reflects the segregation of neuronal populations into functionally related groups^33-35^. Here, modularity was used to quantify functional segregation among neuronal populations within FC networks derived from 2P calcium imaging. Community detection was performed using the Leiden algorithm adapted for signed networks, and modularity (Q) was computed at a fixed resolution parameter (γ = 1) across multiple iterations to ensure stability^36, 37^. Modularity exhibited clear state-dependent differences across all network types. Anesthesia consistently showed the highest modular organization (Q = 0.333 ± 0.005 for positive networks; 0.130 ± 0.017 for negative networks; 0.268 ± 0.036 for combined networks), whereas motion exhibited the lowest modularity (0.258 ± 0.014; 0.086 ± 0.005; and 0.218 ± 0.029, respectively). No-motion networks showed intermediate values. Statistical comparisons confirmed significant differences between anesthesia and motion across all network types (paired t-test, p < 0.05; Fig. 3a), while no-motion did not differ significantly from either condition in positive or combined networks (p > 0.05).

**Figure 3.**
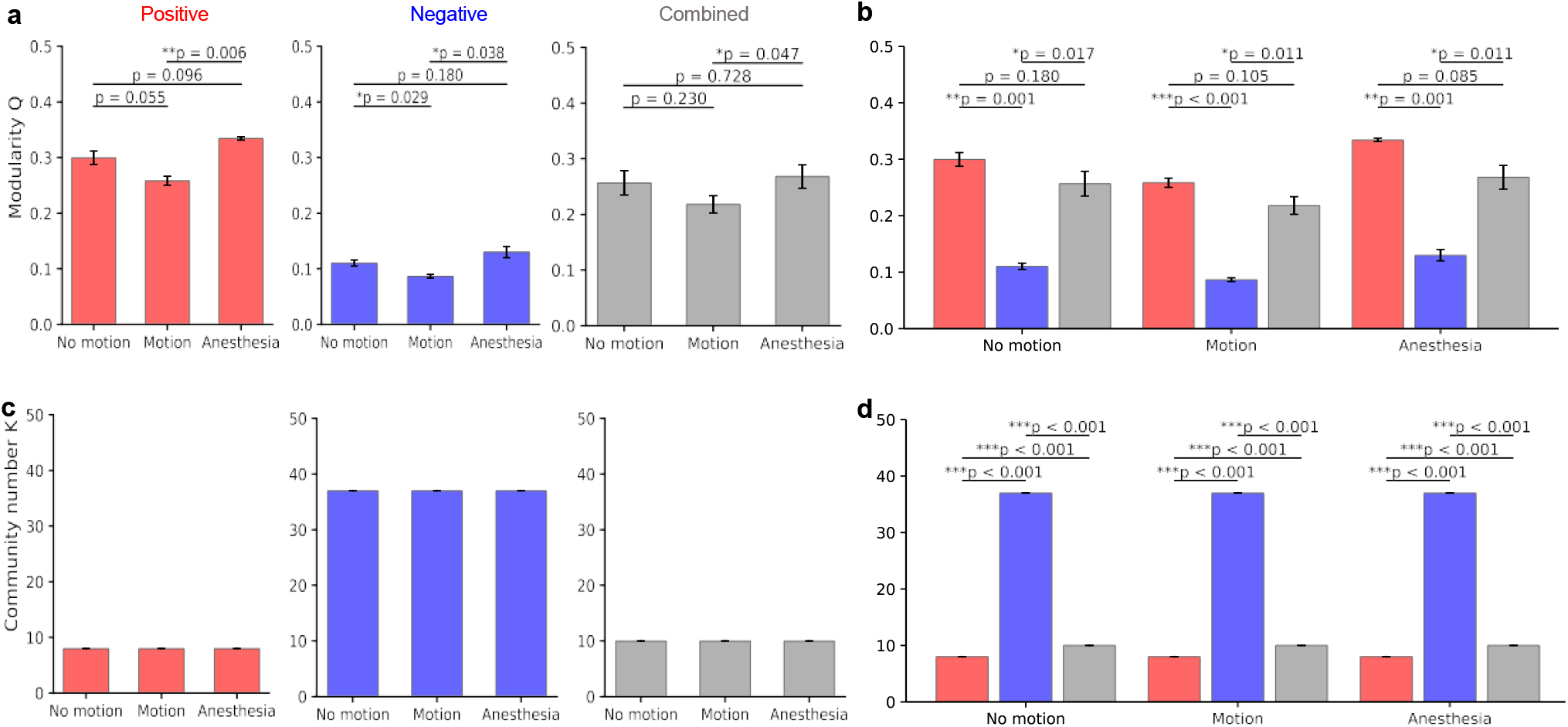
Modularity and community structures of state-dependent networks. (a) Comparison of modularity index *Q* in positive, negative and combined networks. Anesthesia consistently exhibits the highest degree of segregation (*Q*) across all network types compared to the relatively randomized construction of the motion state. (b) Effect of network sign on modularity; the inclusion of negative edges significantly reduces the modularity index (*Q*) of the positive-only backbone. (c) State-wise comparison of community numbers (*K*). The total number of detected communities remains identical within each network type. (d) Community across network types, where negative connectivity results in a significantly higher number of small, fragmented communities. Statistical significance was determined via paired *t*-tests for modularity *Q* and Wilcoxon rank-sum tests for community number *K*; error bars represent SEM; ^*^*p <* 0.05,

Modularity also differed across network sign types. Negative networks exhibited significantly lower modularity compared to both positive and combined networks (paired t-test, p < 0.05). Inclusion of negative edges in combined networks resulted in modest reductions in modularity relative to positive networks, although these differences did not reach statistical significance across states (p > 0.05). A complementary simulation analysis further demonstrated that incorporation of negative associations reduces modular structure and increases variability in community detection. Together, these findings demonstrate that negative connectivity reduces modular segregation in mesoscale functional networks. Community number (K) was derived from Leiden partitions following modularity optimization. The number of detected communities remained stable across behavioral states within each network type (Fig. 3c). Positive networks consistently yielded approximately 8 communities, combined networks approximately 10 communities, and negative networks approximately 37 communities. No significant state-wise differences were observed within each network construction. Across network types, however, community number differed significantly. Negative networks produced substantially higher numbers of communities compared to both positive and combined networks (Wilcoxon rank-sum test, p < 0.001; Fig. 3d), while combined networks also showed higher community counts than positive-only networks (p < 0.001).

In summary, modular organization exhibited both state-dependent and network-type–dependent properties. Modularity followed a consistent trend across states (anesthesia > no-motion > motion), with the strongest community segregation observed under anesthesia and the weakest during motion. Network sign also influenced modular organization: negative networks showed markedly reduced modularity and increased community fragmentation relative to positive and combined networks. Together, these findings demonstrate that both brain state and edge composition shape the modular architecture of mesoscale functional connectivity networks.

### 2.2 Small-world topology

To further characterize global network organization, we next quantified small-world topology of mesoscale functional connectivity networks. Small-world networks are characterized by high local clustering together with short characteristic path lengths, enabling an efficient balance between local specialization and global communication^38-41^. Small-world properties of validated functional connectivity networks were quantified using the Humphries-Gurney index σ, defined as the ratio of normalized clustering coefficient (*C/C*_null_) to normalized characteristic path length (*L/L*_null_)^42^. Degree-preserving randomization was used to generate null networks by conserving the original degree sequence while randomizing edge connections, allowing comparison against equivalent random network topologies.

Small-worldness exhibited clear state-dependent differences across network types. Positive and combined networks showed small-world organization in all states (σ > 1; Fig. 4a). Anesthesia displayed the strongest small-world topology (σ = 3.18 ± 0.26 for positive networks and σ = 2.76 ± 0.04 for combined networks), whereas motion exhibited the lowest σ values, with no-motion occupying an intermediate position. Statistical comparisons confirmed significant differences between anesthesia and the other two states across network types (paired t-test, p < 0.05). In contrast, negative networks consistently showed σ values below 1 (σ = 0.56 ± 0.03 under anesthesia), indicating the absence of small-world organization in negative connectivity patterns (Fig. 4b).

**Figure 4.**
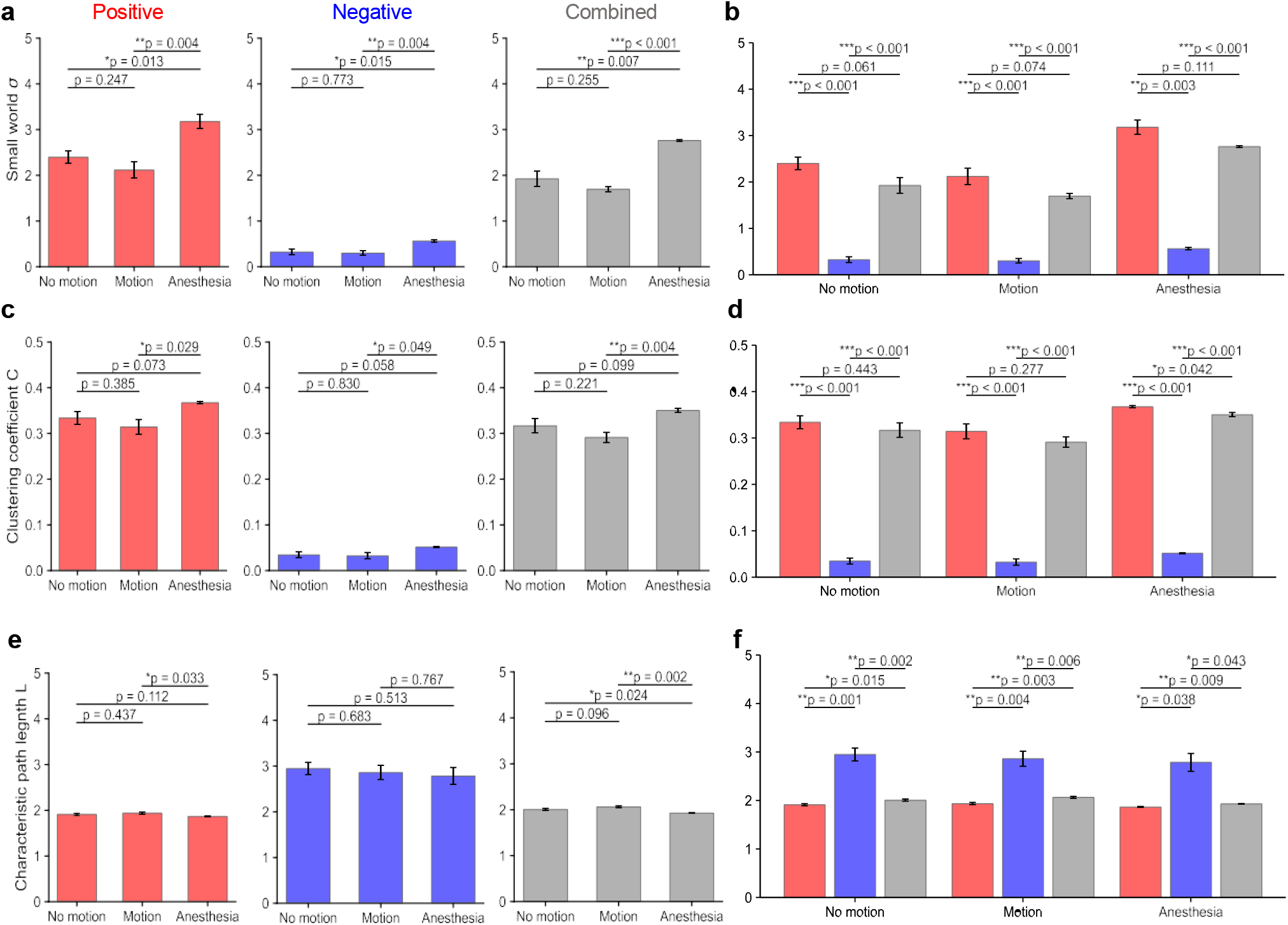
Small-worldness features of state-dependent networks. (a) Comparison of small world index *σ* in positive, negative and combined networks. (b) Effect of network sign on small-worldness. Negative-only networks showed significantly reduced *σ* values below 1, indicating lack of small-world organization. (c-d) Weighted clustering coefficient (*C*) across states and network types. Anesthesia consistently demonstrated the strongest local clustering, with motion showing the lowest values. Negative-only networks exhibited markedly reduced clustering. (e-f) Characteristic path length (*L*) across states and network types. Anesthesia exhibited shorter path lengths than motion in positive and combined networks, indicating greater global accessibility. Negative-only networks showed prolonged path lengths, reflecting inefficient connectivity. Statistical significance was determined via paired *t*-tests; error bars represent SEM; ^*^*p <* 0.05, ^**^*p <* 0.01, ^***^*p <* 0.001.

Network sign also influenced small-world topology. Negative networks exhibited significantly lower σ values than both positive and combined networks (paired t-test, p < 0.001). Inclusion of negative edges in combined networks reduced σ relative to positive networks, although these reductions did not reach statistical significance across all states (p > 0.05). A complementary simulation analysis (Fig. S2) further demonstrated that incorporation of negative associations reduces clustering and increases characteristic path length, resulting in attenuation of small-world organization.

To further examine the structural basis of these differences, the two components of small-worldness-clustering coefficient (C) and characteristic path length (L)—were analyzed separately. Across network types, anesthesia exhibited the highest clustering coefficients (0.367 ± 0.004 for positive, 0.051 ± 0.020 for negative, and 0.350 ± 0.007 for combined networks), followed by no-motion and motion (lowest). Differences between anesthesia and motion were significant across all network types (paired t-test, p < 0.05; Fig. 4c), indicating stronger local connectivity under anesthesia. Characteristic path lengths showed the opposite trend. Anesthesia networks displayed the shortest path lengths (1.93 ± 0.05 for positive, 2.86 ± 0.34 for negative, and 2.06 ± 0.04 for combined networks), whereas motion networks exhibited the longest paths (Fig. 4e). Differences between anesthesia and motion were significant for positive (p = 0.0033) and combined networks (p = 0.002).

Across all states, negative networks exhibited both lower clustering and longer characteristic path lengths than positive and combined networks (paired t-test, p < 0.05; Figs. 4d,f). Inclusion of negative edges in combined networks similarly reduced clustering and increased path length relative to positive-only networks. These effects were most evident under anesthesia, where the proportion of negative edges was higher.

In summary, small-world topology varied systematically across brain states. Anesthesia exhibited the strongest small-world organization, characterized by increased clustering and shorter path lengths. Motion networks showed reduced clustering and longer paths, resulting in weaker small-world structure. Small-world architecture was primarily supported by positive connectivity, whereas negative associations reduced clustering and increased path length, leading to diminished small-world organization.

### 2.3 Hub identification and features comparisons

In network theory, hubs refer to nodes with disproportionately high centrality relative to the rest of the network^43-45^. In mesoscale networks derived from 2P calcium imaging of neuronal population, hubs correspond to neurons occupying highly central positions within the FC network. To identify highly influential neurons within functional connectivity networks, hub detection was based on nodal centrality measures computed from validated networks. Three complementary centrality metrics were used to capture nodal importance from different perspectives: degree centrality, reflecting local connectivity; eigenvector centrality, representing influence within highly connected neighborhoods; and betweenness centrality, quantifying participation in shortest communication paths^46-48^. For each state and network type, hubs were defined as the intersection of the top 10% of nodes ranked by each metric. Examples of hub neurons across the three states are shown in Figs. 5a-c for positive (red), negative (blue), and combined (gray) networks. Hub prevalence was quantified using the hub ratio, defined as the number of hubs divided by the total number of nodes. Across network types, motion exhibited the highest hub ratios (4.39 ± 1.24% for positive networks, 7.67 ± 0.35% for negative networks, and 6.77 ± 0.76% for combined networks), although differences between behavioral states did not reach statistical significance (paired t-test, p > 0.05; Fig. 5d). Anesthesia showed intermediate hub ratios in positive and combined networks despite having the smallest network scales in terms of node and edge counts.

**Figure 5.**
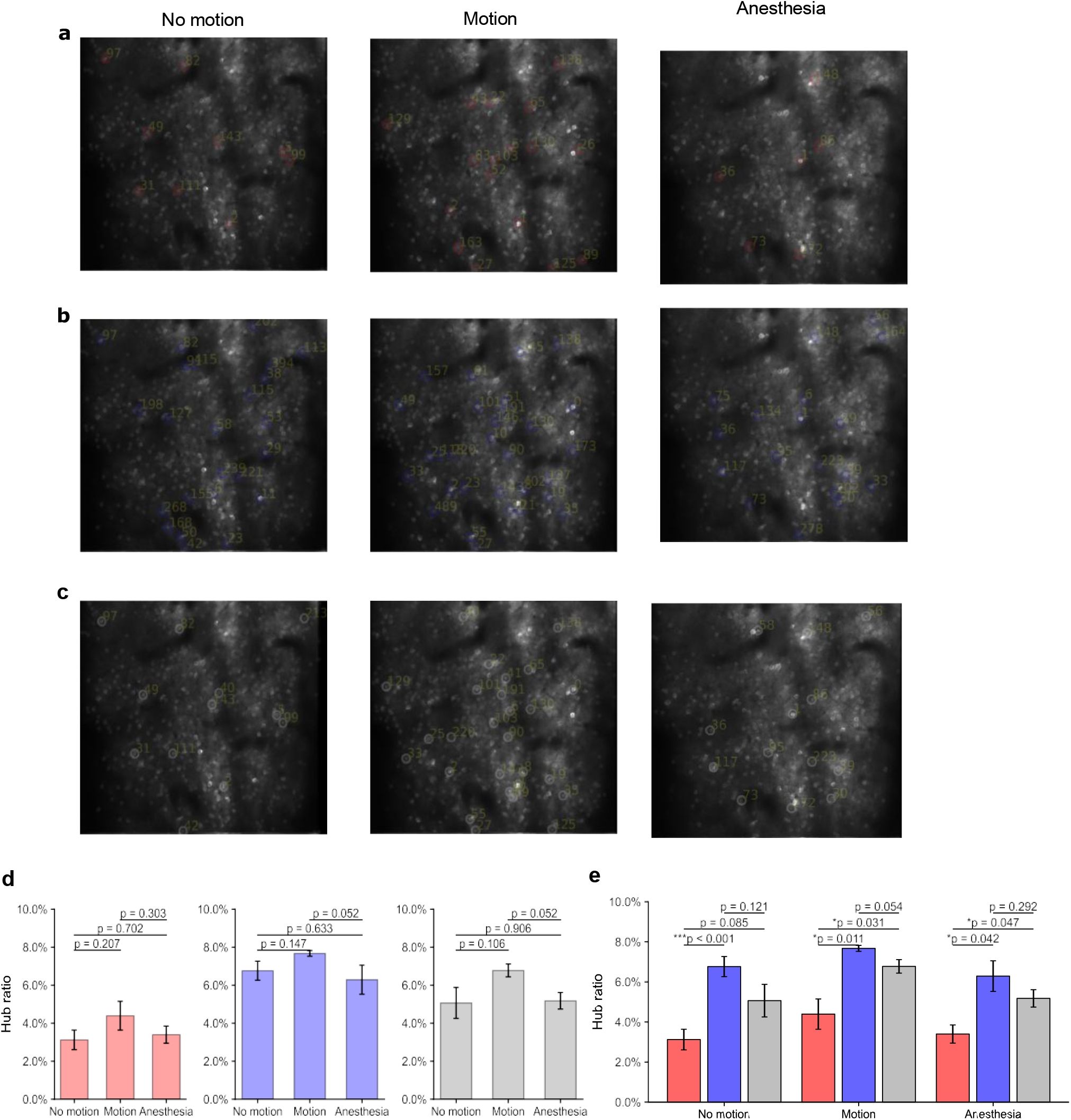
Hub identification and prevalence across behavioral states and network types. (a–c) Examples of hub neurons identified in positive (red), negative (blue), and combined (gray) functional connectivity networks during no motion, motion, and anesthesia states. Hubs were defined as the intersection of the top 10% nodes ranked by degree, eigenvector, and betweenness centralities. (d) State-wise comparison of hub ratios (number of hubs divided by total node count) across network types. (e) Network-type comparison of hub ratios across behavioral states. Negative-only networks exhibit significantly higher hub prevalence under the multi-metric intersection criterion. Statistical significance was determined via paired *t*-tests; error bars represent SEM; ^*^*p <* 0.05, ^**^*p <* 0.01, ^***^*p <* 0.001.

Hub prevalence also differed across network types. Negative networks exhibited significantly higher hub ratios than positive and combined networks (Fig. 5e), indicating greater overlap among the centrality metrics used for hub detection. In positive networks, degree and eigenvector centralities primarily reflected local connectivity structure, whereas betweenness centrality identified nodes participating in shortest communication paths. As a result, fewer nodes simultaneously satisfied all three centrality criteria. In contrast, negative networks showed stronger agreement among the centrality measures, leading to greater multi-metric hub overlap. A complementary simulation analysis supported this observation (Fig. S3, Supplementary File).

To further characterize hub organization across behavioral states and network types, several nodal properties were examined, including mean ΔF/F activity, hub degree, connection strength, and spatial distances between hubs. For neuronal activity profiles, anesthesia hubs consistently exhibited the lowest ΔF/F amplitudes (0.65 ± 0.19 in positive networks, 0.77 ± 0.10 in negative networks, and 0.70 ± 0.14 in combined networks), whereas motion hubs showed the highest activity levels (paired t-test, p < 0.05; Fig. 6a). Differences among network types were not statistically significant (p > 0.05; Fig. 6b). These results indicate reduced hub-associated activity under anesthesia and increased activity during motion. Hub degree showed a distinct state-dependent pattern. Across all network types, anesthesia networks exhibited the highest hub degrees (87.55 ± 7.38 for positive, 50.38 ± 4.20 for negative, and 123.25 ± 14.51 for combined networks), whereas motion networks consistently showed the lowest hub degrees (Fig. 6c). Differences between anesthesia and motion reached statistical significance in negative (p = 0.023) and combined networks (p = 0.027; paired t-test). No-motion networks showed intermediate hub degrees across all network types. Across network types, negative networks exhibited significantly lower hub degrees than positive and combined networks (Fig. 6d), consistent with the overall lower edge density observed in negative connectivity networks. Hub strength (mean connection weight) followed a similar pattern. Anesthesia hubs showed higher connection strengths in all network types (0.168 ± 0.006 in positive networks, 0.070 ± 0.013 in negative networks, and 0.159 ± 0.004 in combined networks), whereas motion and no-motion hubs exhibited lower and intermediate strengths, respectively. Differences between anesthesia and motion were significant in positive (p = 0.002) and combined networks (p = 0.027; paired t-test; Fig. 6e).

**Figure 6.**
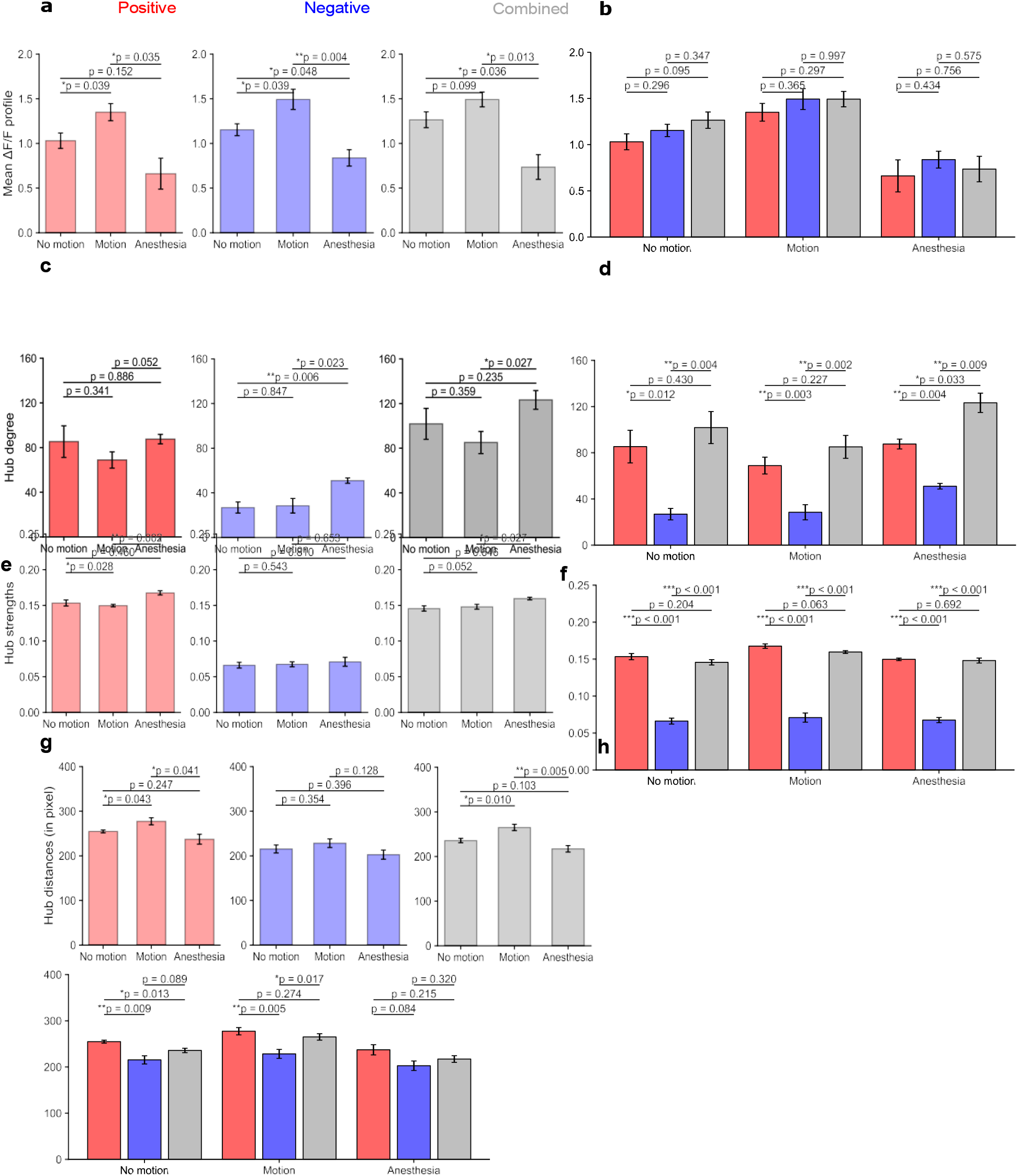
Hub features comparison. (a–b) Mean Δ*F/F* activity of hub neurons across behavioral states (a) and network types (b). (c-d) Hub degree (number of edge connections) across behavior states (c) and network types (d). (e-f) Mean hub edge strength (average edge weight) across behavior states (e) and network types. (g–h) Mean pairwise spatial distance between hub neurons across behavior states (g) and network types (h). Anesthesia is characterized by reduced hub activity but elevated hub degree and connection strength, whereas motion exhibits higher activity with comparatively weaker structural embedding. Negative-only networks display reduced hub degree and strength relative to positive and combined networks. Statistical significance was determined via paired *t*-tests; error bars represent SEM; * *p <* 0.05, ^**^*p <* 0.01, ^***^*p <* 0.001.

Across network types, negative networks showed significantly lower hub strengths compared with positive and combined networks (Fig. 6f). Inclusion of negative edges in combined networks slightly reduced hub strengths relative to positive-only networks, although these reductions were not statistically significant (p > 0.05). Spatial organization of hubs also differed across states. Hub–hub distances were shortest under anesthesia and longest during motion across all network types (Fig. 6g), with significant differences observed in positive (p = 0.041) and combined networks (p = 0.005). Across network types, negative hubs generally showed shorter spatial separations than positive and combined hubs, although significance varied by state (Fig. 6h). Reduced hub distances under anesthesia were consistent with the smaller network scales and increased clustering observed in this state.

In summary, hub organization varied across both brain states and network types. Motion networks exhibited the highest hub ratios, whereas anesthesia networks maintained intermediate hub prevalence despite smaller network scales. Hub properties also differed across states: anesthesia networks showed hubs with higher connectivity (degree and strength) but lower neuronal activity, whereas motion networks exhibited higher hub activity with comparatively weaker connectivity. Across network types, positive-network hubs maintained stronger connection weights than negative-network hubs, while negative hubs, although more prevalent under the intersection criterion, were characterized by weaker individual connections.

## Discussion

Mesoscale neuronal population networks represent an intermediate organizational level linking single-neuron activity to large-scale brain networks^1, 11^. Using in vivo 2P calcium imaging combined with graph-theoretical analysis, we examined the topological organization of these networks across behavioral states. Motion networks exhibited the largest FC architectures, whereas anesthesia networks showed reduced network scales together with stronger modular segregation and more pronounced small-world topology. Hub analyses revealed additional state-dependent differences: anesthesia networks showed stronger hub connectivity despite reduced neuronal activity, whereas motion networks exhibited higher hub activity with comparatively weaker connectivity.

These findings are consistent with the view of the brain as a complex system composed of interacting networks across multiple spatiotemporal scales^8, 9^. Within this framework, network topology reflects the balance between functional segregation and integration. The increased modular segregation and stronger small-world organization observed under anesthesia indicate a more locally structured mesoscale configuration. In contrast, motion was associated with reduced modularity and weaker small-world topology, reflecting a shift toward more distributed network organization. Importantly, these patterns were evident at the level of neuronal population networks measured with 2P calcium imaging, indicating that state-dependent reorganization is present at the mesoscale level of cortical networks.

Network sign also contributed to the observed mesoscale organization. Across behavioral states, networks constructed from negative associations consistently showed reduced modularity, lower clustering coefficients, longer characteristic path lengths, and small-world indices below one. In contrast, positive FC networks preserved modular community structure and small-world topology. When negative associations were incorporated into combined networks, modest reductions in modularity and small-worldness were also observed. These results indicate that the balance between positive and negative correlations influences the global organization of mesoscale FC networks.

Hub organization also differed across states and network types. Anesthesia networks exhibited hubs with higher connectivity degree and strength despite reduced neuronal activity, whereas motion networks showed higher hub activity but weaker connectivity structure. Across network types, positive-network hubs maintained stronger connection weights than negative-network hubs, while negative hubs were more prevalent under the multi-metric intersection criterion but characterized by weaker individual connections. These findings indicate that hub organization in mesoscale networks depends on both brain state and network composition.

Several limitations should be considered. FC in this study was derived from pairwise correlations of calcium activity and therefore reflects statistical associations rather than direct synaptic interactions. In addition, 2P imaging samples neuronal populations within a restricted cortical field of view, limiting inference about network organization at larger spatial scales. Future work combining mesoscale calcium imaging with larger-scale recording approaches will help link neuronal population dynamics to broader brain network organization.

Overall, these findings show that mesoscale neuronal population networks exhibit structured and state-dependent topological organization. Combining 2P calcium imaging with graph-theoretical analysis provides a framework for examining mesoscale cortical dynamics across behavioral states, in normal brain function and brain disorders.

## Funding

This study was supported by the National Science Foundation (NSF grant 2242771) and GM109098-09S1 Supplement from NIH Centers of Biomedical Research Excellence (COBRE).

## Author Contribution

S.L. conceived and designed the study. G.P. and S.L. developed the methodology. N.S. performed the experiments. G.P. analyzed the network data. G.P. and S.L. wrote and edited the manuscript.

## Conflict of Interest

The authors declare no competing interests.

## Data and code availability

All data generated in this study will be made available from the corresponding author upon reasonable request, and all analysis code is publicly available at LatifiLab GitHub (https://github.com/LatifiLab/neural_decoding).

## Notes

### Competing Interest Statement

The authors have declared no competing interest.

